## Supplementary Information for "Cytonemes coordinate asymmetric signaling and organization in the *Drosophila* muscle progenitor niche"

**This file includes**

Supplementary Figures 1-7

Supplementary Tables 1-3

References

**Supplementary Figure 1. Characterization of AMP localizations in the wing disc niche.**

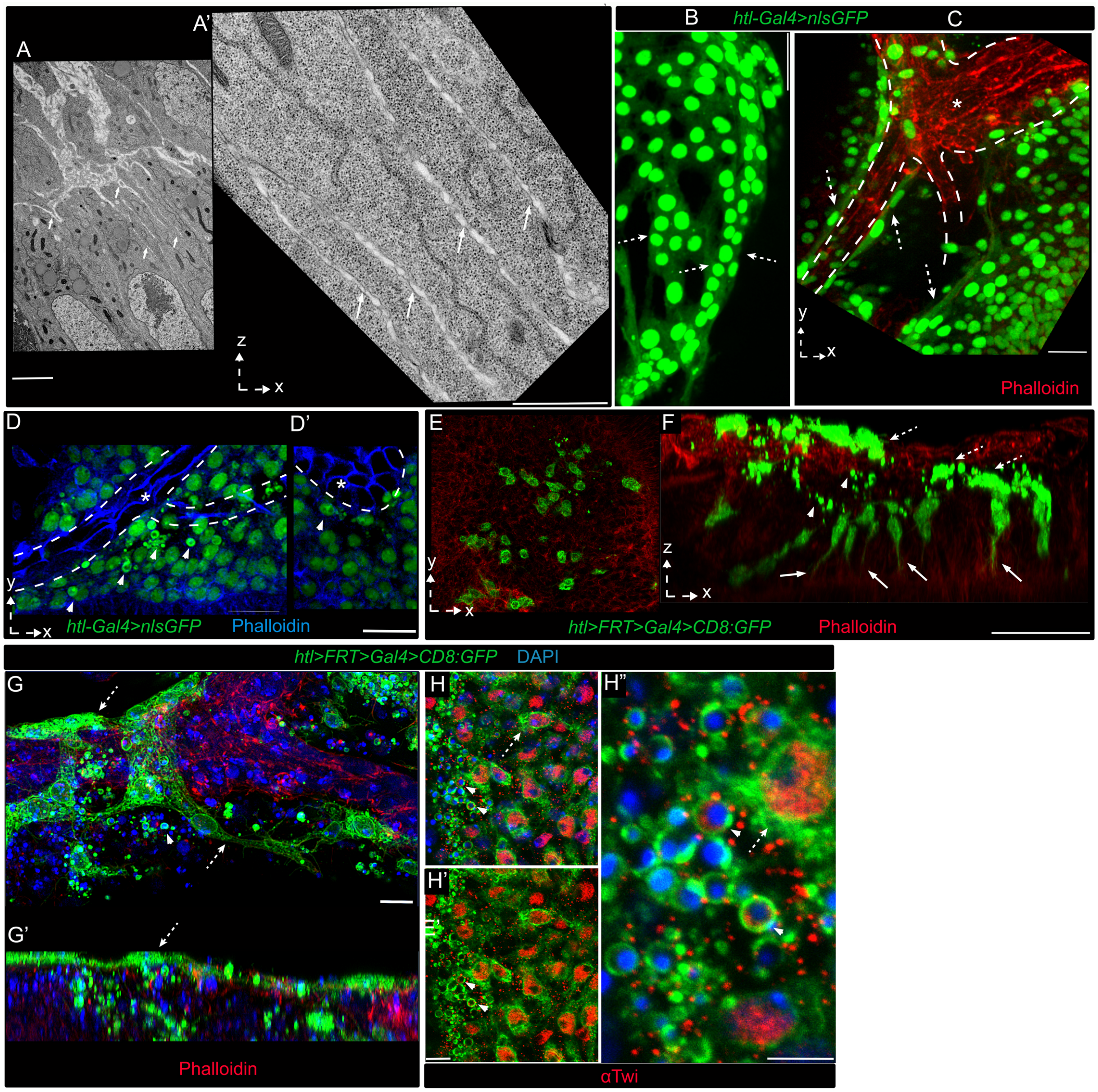


**A-A'** TEM sections of a $w^{1118}$ wing disc showing cytoneme-like AMP projections within the intercellular space of the wing disc epithelium. **B-D'** XY sections showing diverse morphology of distal layer AMPs; arrowhead, small non-polar cells; dashed arrow, multinucleated elongated cells in juxtaposition to the trachea (*, dashed lines, phalloidin stained) (also see Fig.1I-K). **E-H"** Images of a wing disc harboring CD8:GFP-marked AMP clones; E, F, a single optical XY section; F, YZ view of a wing disc showing orthogonal (arrows) and lateral (dashed arrows) orientation of cells and cytonemes, occurring exclusively in proximal (arrow) and distal (dashed arrow) clones, respectively, relative to the disc plane; G-G', Clones of distal cells showing diverse morphologies, cell-cell adhesion, multi-nucleated assembly; red, phalloidin; H-H", Twi-immunostained tissues showing small spherical cells, large elongated cells; dashed arrow, elongated cells; arrowhead, spherical cells. Scale bars: 20μm; 5μm (A); 2μm (A'); 20μm (B-F); 5μm (G-H").

**Supplementary Figure 2. Characterization of AMP cytonemes.**

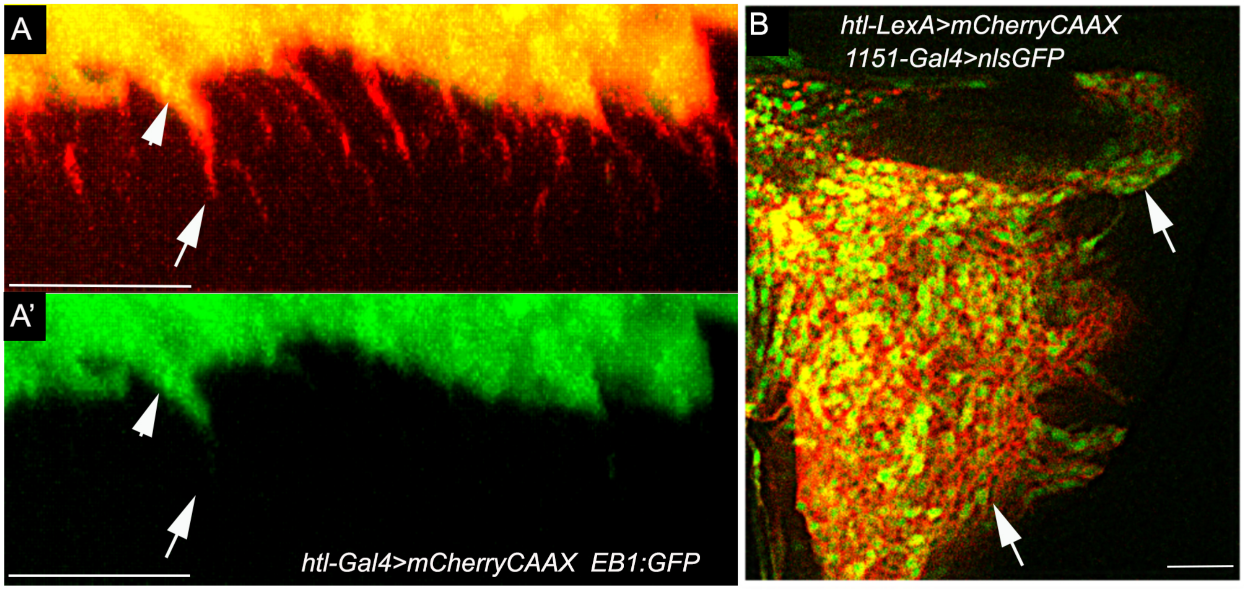


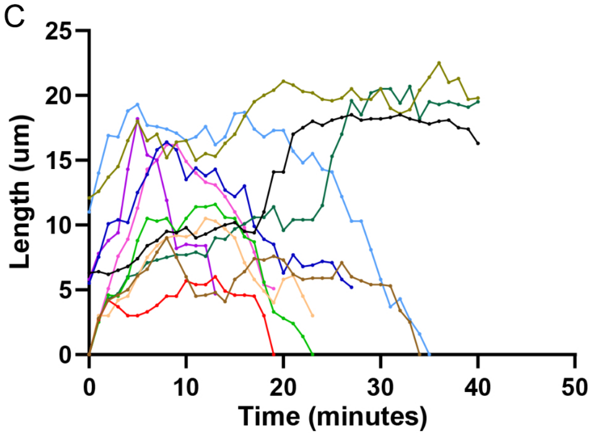


D

| *htl-LexA>CD2:GFP* | | # Cytonemes | # of discs |
|---|---|---|---|
| Average Maximum extension (μm) | 14.9 ± 4.4 | 19 | 4 |
| Average Projection velocity (μm /min) | 1.2 ± 0.8 | 19 | 4 |
| Average Retraction velocity (μm /min) | 0.8 ±0.4 | 19 | 4 |
| Average Lifetime (min) | 25.07 ±11.3 | 105 | 4 |

**A-A'** mCherryCAAX-marked AMPs expressing EB1:GFP lack GFP signal at the growing tips of orthogonal cytonemes; Arrowheads, cytonemes base; arrows, cytonemes shaft, and tip. **B** Highly specific pan-AMP expression pattern of *htl-LexA* binary transcription driver as verified by the overlap of expression pattern (arrow) over the known *1151-Gal4* pan-AMP driver. **C** Kymograph showing the dynamics of *htl-LexA>LexO*-CD2:GFP marked AMP cytonemes; each line graph indicates tracking of a single filopodium (4 discs). **D.** Analyses of cytoneme dynamics; average values ± SD were derived from >19 cytonemes in four *ex vivo* cultured wing discs as indicated. Source data are provided as a Source Data file. Scale bars: 20μm.

**Supplementary Figure 3. Generation and expression of *pyr-Gal4*.**

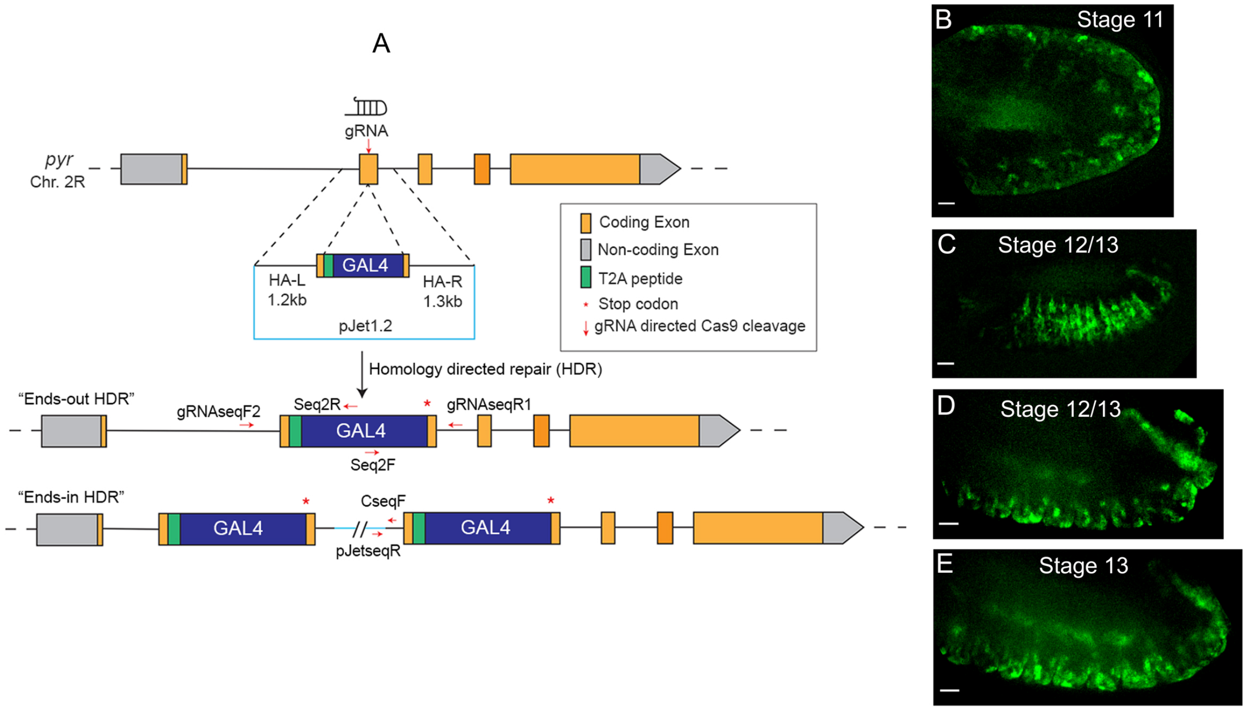


**A** Schematic illustration of CRISPR/Cas9-based genome editing to generate *pyr-Gal4* transgenic *Drosophila*; indicated primers were used to screen "ends-out" HDR lines (See Materials and Methods). **B-E** Different stages of embryos expressing CD8:GFP under *pyr-Gal4*; *pyr-Gal4* expression patterns matched previously published *pyr* mRNA *in situ* hybridization patterns, e.g., embryonic ectoderm near pericardial cell precursors (B), lateral view of segmental epithelial stripes (C), and ventral epithelial expression near proctodeum and stomodeum (D,E). Scale bars: 50μm.

**Supplementary Figure 4. Niche-specific signaling of Pyr:GFP and Ths:GFP.**

A

| Genotype | # of discs | AMP location | # of AMPs | Av. thickness of ad-epithelial layer (μm) | # of layers |
|---|---|---|---|---|---|
| *htl>nls:mCherry* | 3 | Hinge | 18.7±4.0 | 5.7±0.4 | 1.7 ±0.6 |
| | | Notum | 31.3±3.8 | 28.7± 4.6 | 4.7 ±0.6 |
| *pyr>Pyr:GFP, htl>nls:mCherry* | 3 | Hinge | 35.0±3.6 *** | 15.0±1.3 *** | 3.7 ±0.6 *** |
| | | Notum area | 48.0±11 | 45±2.2 ** | 6.7 ±0.6 |
| *ths>Ths:GFP, htl>nls:mCherry* | 3 | Hinge | 18.0±4.4 | 6.8±1.4 | 2.0 ±0.0 |
| | | Notum | 102.3±19.6 *** | 59.4±7.5 *** | 13.3±1.5 *** |

**, p<0.05; ***, p<0.01 (t-test compared to *htl-LexA*>nls:mCherry).

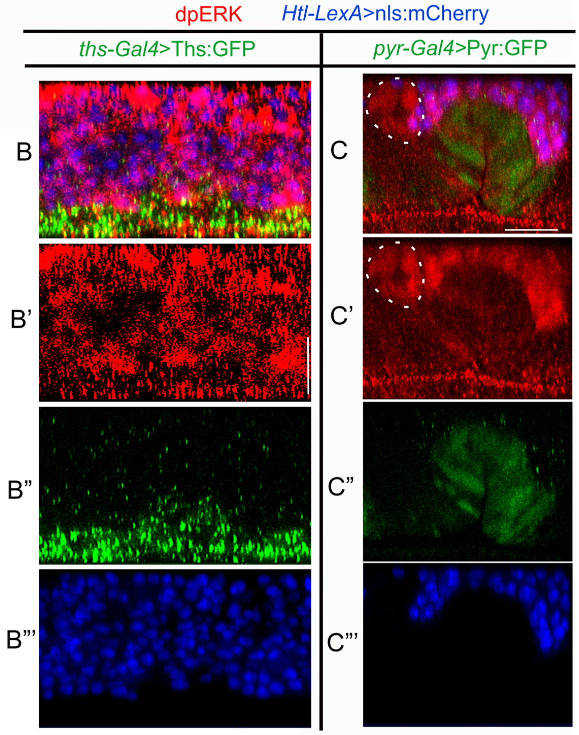


**A-C'''** Wing discs expressing Pyr:GFP and Ths:GFP under *pyr-Gal4* and *ths-Gal4*, respectively. **A** Niche-specific change in the AMP pool size and stratified organization due to the Pyr:GFP and Ths:GFP expression from their respective sources; hinge, *pyr* source; notum, *ths*-source; Average values ± SD shown; **, p<0.05; ***, p<0.01 (unpaired two-tailed t-test). **B-C'''** Activation of dpERK (red) in all signal-receiving AMPs (blue); dashed circle, ASP. Scale bars: 20μm. Source data are provided as a Source Data file.

**Supplementary Figure 5. AMP homing on ectopic Pyr:GFP and Ths:GFP-expressing source.**

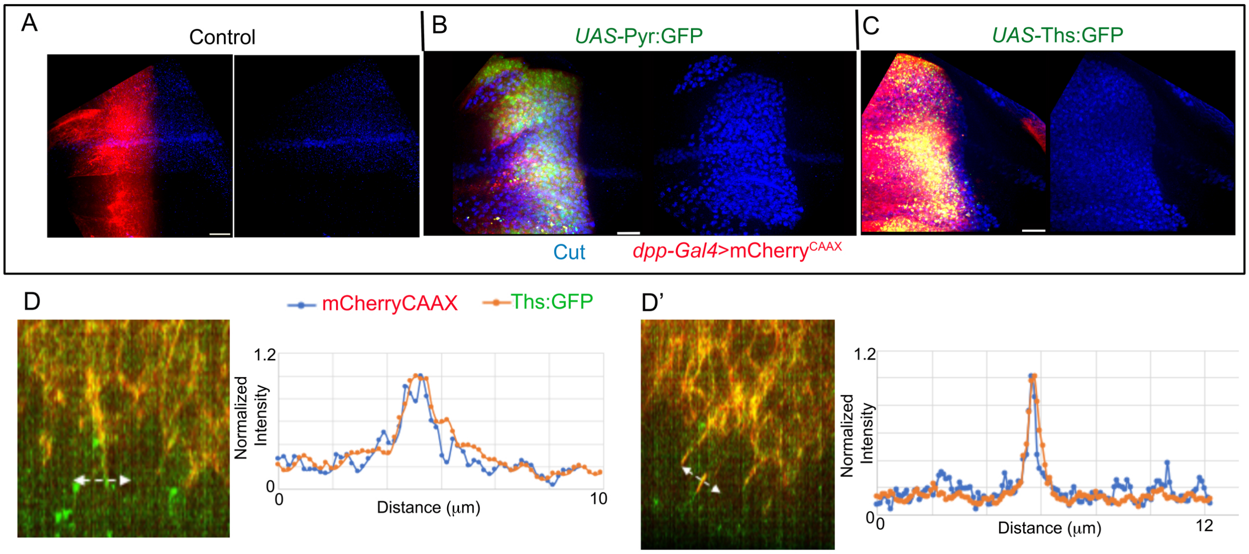


**A-C** The homing of AMPs (marked with αCut) over the disc *dpp*-source ectopically expressing of Pyr:GFP (B) or Ths:GFP (C); A, control disc region (*dpp-Gal4, UAS-mCherryCAAX/+*). **D-D'** Raw quantitation data from single XZ confocal slices from wing discs to assess colocalization of Ths:GFP produced from the *dpp* source in the disc pouch and mCherryCAAX-marked AMP cytonemes that invade into the ectopic niche. Genotypes: *dpp-Gal4/LexO-mCherryCAAX; htl-LexA/UAS-Ths:GFP* (D, D'). Source data are provided as a Source Data file. Scale bars: 20µm.

**Supplementary Figure 6. AMP cytonemes in the Wg-expressing disc source.**

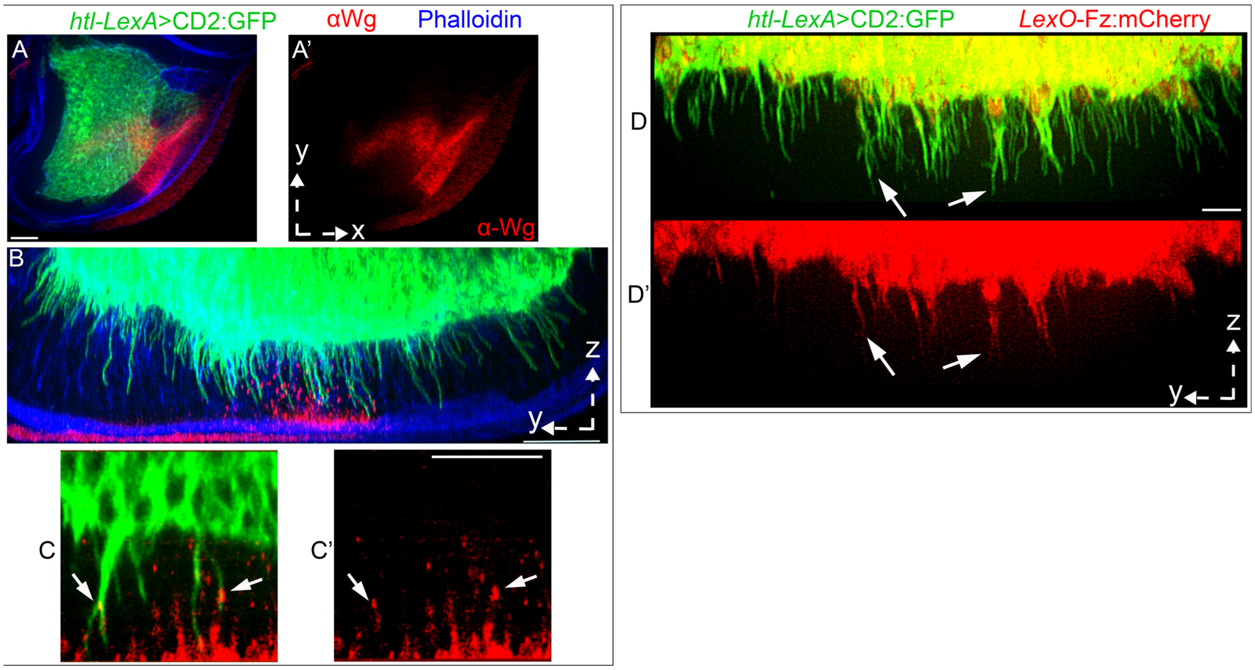


**A,A'** CD2:GFP-marked AMPs over the Wg-expressing (αWg; red) zone in the wing disc. **B,C** A group of AMP cytonemes that occupied the *ths*-expressing wing disc notum also receive Wg (αWg; red; arrow in C). **D, D'** CD2:GFP-marked AMPs expressing *LexO*-Fz:mCherry under *htl-LexA* showed Fz:Cherry-containing cytonemes specifically within the Wg-expressing zone (arrows). Scale bars: 20μm.

**Supplementary Figure 7. Non-autonomous effects of cytoneme-deficient AMPs on the wing disc.**

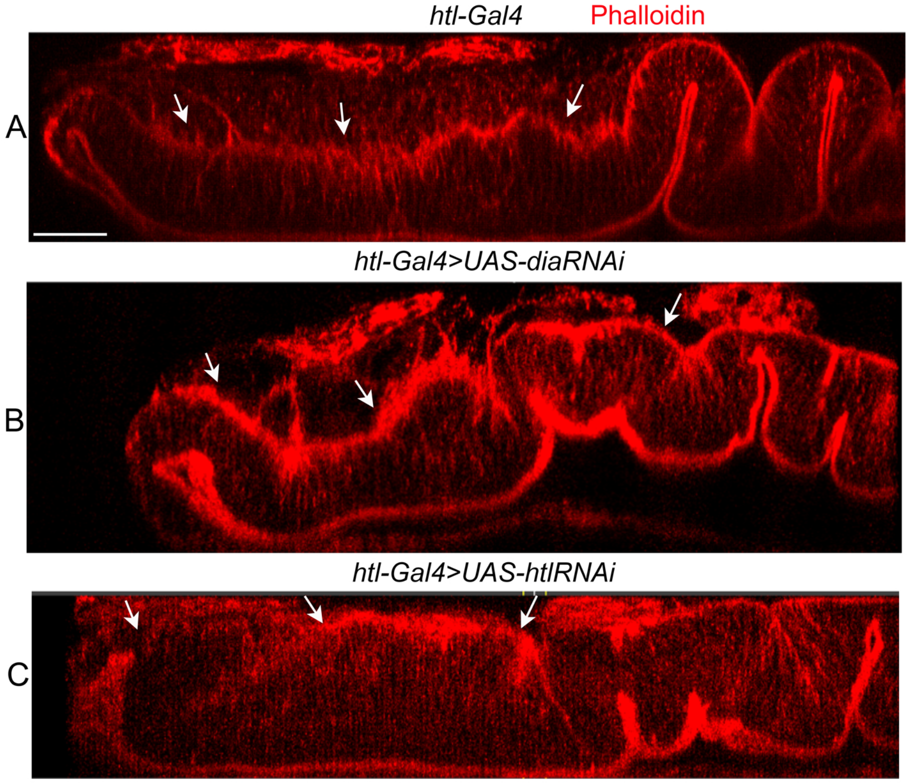


**A-C** YZ sections of wing imaginal discs, comparing identical region of discs derived from WT control (*htl-Gal4 X* $w^-$) and mutant flies that expressed either *dia-i* (B) or *htl-i* (C) under *htl-Gal4* (*htl-Gal4 X UAS-RNA-i*); red, phalloidin staining; arrows, basal surface of wing disc. Note that *dia-i* expressing syncytial AMPs caused abnormal disc morphology, probably to accommodate large-sized AMPs (also see Figure 4H-I). Similarly, the loss of disc-specific polarity of *htl-i*-expressing AMPs/AMP cytonemes caused smoothened disc surface and reductions in wing disc folds and actin-rich projections (also see Figure 5G-J). Scale bars: 20μm.

### SUPPLEMENTARY TABLES

#### Supplementary Table 1. Polarized AMP morphology and organization in niche

| | Genotype | # of discs | AMP layer | Av. major axis length orthogonal or oblique to the disc (μm) ± SD | Av. major axis length parallel to the disc (μm) ±SD | # of AMPs | Angle between major axes of AMPs and disc (av. deg.) ± SD | Av. thickness of the ad-epithelial layer (μm) ± SD | # of layers ± SD |
|---|---|---|---|---|---|---|---|---|---|
| EM | $w^-$ | 2 * | p | 7.1±1.5 | 3.1±0.6 | 16 | 81.9±7.8 | 21.5± 1.5 | 4.0 ±0.0 |
| | | | d | 2.3±1.1 | 8.7±2.6 | 11 | 11.7±4.8 | | |
| Light microscopy | *htl-Gal4> nls:GFP* | 5 | p | 5.5±1.2 | 2.9±0.6 | 125 | 83.4±5.5 | 26.0±1.2 | 4.5±0.7 |
| | | | $p^{-1}$ | 4.7±1.1 | 3.7±0.9 | 119 | 77.2±9.5 | | |
| | | | $d^{-1}$ | 3.9±1.1 | 4.3± 0.9 | 84 | 21.0±13.1 | | |
| | | | d | 3.2±1.1 | 5.2±1.1 | 58 | 8.8±6.0 | | |
| | *htl-Gal4> nls:GFP, htl-i* | 3 | p | 3.1±0.8 ** | 6.2±1.9 ** | 33 | 6.0±4.3 ** | 5.9±1.3 ** | 1.0±0.0 *** |

| *htl-Gal4> nls:GFP, dia-i* | # of discs | Av. diameter of myoblasts (μm) | Average thickness of myoblasts (μm) | Average nuclei per chamber | Average number of chambers |
|---|---|---|---|---|---|
| | 5 | 17.5 ±10.9 | 50.5 ±12.6 | 6.5 ±1.2 | 6.4 ±1.1 |

**Note:** p, proximal. d, distal. $p^{-1}$, one layer above p relative to the disc. $d^{-1}$, one layer below d relative to the disc. Source data are provided as a Source Data file.

*, 16 TEM sections from 2 wing discs.

**, p <0.0001, unpaired two-tailed t-test compared to *htl>nls:GFP*.

***, p = 0.0002, unpaired two-tailed t-test compared to *htl>nls:GFP*.

**Supplementary Table 2. Clonal analyses of AMP and AMP cytonemes**

| Genotype | | # of discs | Clone layer | # Clones * | Av. # disc-directed cytoneme/cell ± SD | Av. # lateral cytoneme/cell ± SD | Av. # of non-polar small cells ± SD | Av. cell angle (av. Deg. ± SD) |
|---|---|---|---|---|---|---|---|---|
| *htl>FRT>Gal4* | *UAS-CD8:GFP* | 3 | p | 41 | 3.2±0.9 | 0.0 | 0 | 83.2±4.7 |
| | | | d | 15 | 0 | 5.7± 1.3 | 39.7± 21.9 | 2.5±3.2 |
| | *Lifeact:GFP* | 3 | p | 25 ** | 2.6±0.9 ** | 0 | 0 | 85.7±5.3 |
| | | | d | 19 | 0 | 5.6± 1.8 | 36.7± 9.5 | 1.1±1.2 |
| | *Lifeact:GFP, dia-i* | 9 | p | 0 ** | 0 ** | ND | 0 | NA |
| | | | d | NA | NA | ND | 71.6± 66.3 | NA |
| | *Lifeact:GFP, htl-i* | 10 | p | 0 ** | 0 ** | ND | 0 | NA |
| | | | d | 24 | 0 | 6.1± 1.7 | 57.3± 37.4 | 0.79±1.2 |

Note: *, excludes non-polar spherical cells. p, proximal. d, distal. **, $p < 0.01$ (One-way ANOVA followed by Tukey HSD) comparing same myoblast layers between control (*htl>FRT>Gal4>Lifeact:GFP*) and *htl-i and dia-i*. Source data are provided as a Source Data file.

### Supplementary Table 3. Reagent list used in this study

| REAGENT or RESOURCE | SOURCE | IDENTIFIER |
|---|---|---|
| Antibodies | | |
| Mouse anti-MAP Kinase, Activated (Diphosphorylated ERK-1&2) (1:250) | Sigma-Aldrich | Cat# M-8159; RRID:AB_477245 |
| Rabbit anti-PH3 (1:2000) | Cell Signaling Technology | Cat# 9701; RRID:AB_331535 |
| Mouse anti-Cut (1:50) | DSHB | Cat# 2B10; RRID:AB_528186 |
| Mouse monoclonal anti-Discs large (1:100) | DSHB | Cat# 4F3 anti-discs large; RRID: AB_528203 |
| Mouse anti-Armadillo (1:100) | DSHB | Cat# N2 7A1; RRID:AB_528089 |
| Mouse anti-Wingless (1:50) | DSHB | Cat# 4D4; RRID:AB_528512 |
| Rat anti-Shotgun (1:50) | DSHB | Cat# DCAD2; RRID:AB_528120 |
| Rabbit anti-Twist (1:2000) | [1] | N/A |
| Rabbit anti-Vestigial (1:200) | [2] | N/A |
| Goat anti-Mouse IgG (H+L), Alexa Fluor 555 | Thermo Fisher Scientific | A21434 |
| Goat anti-Mouse IgG (H+L), Alexa Fluor 647 | Thermo Fisher Scientific | A28181 |
| Goat anti-Rat IgG (H+L), Alexa Fluor 647 | Thermo Fisher Scientific | A21247 |
| Goat anti-Rabbit IgG (H+L), Alexa Fluor 555 | Thermo Fisher Scientific | A21428 |
| Goat anti-Rabbit IgG (H+L), Alexa Fluor 647 | Thermo Fisher Scientific | A21244 |
| Bacterial and Virus Strains | | |
| DH5 Alpha | | |
| Chemicals, Peptides, and Recombinant Proteins | | |
| Phalloidin iFlor 555 | Abcam | Cat# ab176759 |
| Phalloidin iFlor 647 | Abcam | Cat# ab176756 |
| Sodium Cacodylate | Electron Microscopy Sciences | Cat# 12300 |
| Osmium Tetroxide | Electron Microscopy Sciences | Cat# 19140 |
| Potassium Ferricyanide | Sigma-Aldrich | Cat# 702587 |
| Uranyl Acetate | Electron Microscopy Sciences | Cat# 22400 |
| Propylene Oxide | Electron Microscopy Sciences | Cat# 20401 |
| Low Viscosity Resin | Electron Microscopy Sciences | Cat# 14300 |

| | | |
|---|---|---|
| Lead Citrate | Electron Microscopy Sciences | Cat# 17800 |
| Poly-L-lysine | VWR | Cat# 48393241 |
| Critical Commercial Assays | | |
| CloneJET PCR Cloning Kit | Thermo Fisher Scientific | Cat# K1231 |
| Gateway™ LR Clonase™ II Enzyme mix | Thermo Fisher Scientific | Cat# 11791020 |
| Zymoclean Gel DNA Recovery Kit | Zymo Research | Cat# D4007 |
| GeneJET Plasmid Miniprep Kit | ThermoFisher Scientific | Cat# K0502 |
| GeneJET Plasmid Midiprep Kit | ThermoFisher Scientific | Cat #K0481 |
| 2X PCR Premix | Syd Labs | Cat# MB067-EQ2R-L |
| Deposited Data | | |
| Raw data from all the figures | This paper | |
| Experimental Models: Organisms/Strains | | |
| *D. melanogaster*: *UAS-CD8:GFP* | BDSC | 5130 |
| *D. melanogaster*: *UAS-CD8:GFP* | BDSC | 5137 |
| *D. melanogaster*: *UAS-CD8:RFP* | BDSC | 32218 |
| *D. melanogaster*: *UAS-mCherryCAAX* | BDSC | 59021 |
| *D. melanogaster*: *lexO-mCherryCAAX* | [3] | N/A |
| *D. melanogaster*: *lexO-CD2:GFP* | BDSC | 66544 |
| *D. melanogaster*: *UAS-Lifeact:GFP* | BDSC | 57326 |
| *D. melanogaster*: *UAS-Eb1:GFP* | BDSC | 35512 |
| *D. melanogaster*: *UAS-nls:GFP* | BDSC | 4776 |
| *D. melanogaster*: *UAS-nls:mCherry* | BDSC | 38425 |
| *D. melanogaster*: *UAS-Dia-GFP* | BDSC | 56751 |
| *D. melanogaster*: *UAS-ΔDAD-Dia-GFP* | BDSC | 56752 |
| *D. melanogaster*: *LexO-nsyb:*$GFP_{1\text{-}10}$*,* *UAS-CD4:*$GFP_{11}$ | [4] | N/A |
| *D. melanogaster*: *UAS-htl-DN* | BDSC | 5366 |
| *D. melanogaster*: *UAS-htlACT* | BDSC | 5467 |
| *D. melanogaster*: *UAS-pyrRNAi* | BDSC | 63547 |
| *D. melanogaster*: *UAS-diaRNAi* | BDSC | 33424 |
| *D. melanogaster*: *htl-Gal4* | BDSC | 40669 |
| *D. melanogaster: ths-Gal4* | BDSC | 77475 |
| *D. melanogaster*: *{nos-Cas9}ZH-2A* | BDSC | 54591 |
| *D. melanogaster*: *hs-Flp* | BDSC | 6 |
| *D. melanogaster*: $w^{1118}$ | BDSC | 3605 |
| *D. melanogaster*: *htl:*$GFP^{fTRG}$ | VDRC | 318120 |
| *D. melanogaster*: *UAS-htlRNAi* | VDRC | 6692 |
| *D. melanogaster*: *UAS-thsRNAi* | VDRC | 24536 |
| *D. melanogaster: dpp-Gal4* | [5] | N/A |
| *D. melanogaster*: *LexO-Fz:mCherry* | [5] | N/A |
| *D. melanogaster*: *1151-Gal4* | [5] | N/A |
| *D. melanogaster*: *htl-LexA* | This paper | N/A |
| *D. melanogaster*: *pyr-Gal4* | This paper | N/A |
| *D. melanogaster*: *htl>FRT>stop>FRT>Gal4* | This paper | N/A |

| | | |
|---|---|---|
| *D. melanogaster*: *LexO-Htl:mCherry* | This paper | N/A |
| *D. melanogaster: UAS-Ths:GFP* | This paper | N/A |
| *D. melanogaster*: *UAS-Pyr:GFP* | This paper | N/A |
| Oligonucleotides | | |
| Primer for cloning *pyr-Gal4*: AGGACTTATATTATACTGATGGTGAGTTTTTGTCC | This paper | N/A |
| Primer for cloning *pyr-Gal4*: AATTCGAGCTCGGTACCCTTCTGCTATTGATCTGC CAGCG | This paper | N/A |
| Primer for cloning *pyr-Gal4*: GCCCAATGTCGAATTCGGCTCCGGCGAAGG ACGCGGCAGCCTACTGACTTGCGGAGATGTCGAA GAG AACCCTGGCCCTATGAAGCTACTGTCTTCTATC | This paper | N/A |
| Primer for cloning *pyr-Gal4*: CGCCGGAGCCGAATTCGACATTGGGCATGAACTT GTGGAAC | This paper | N/A |
| Primer for cloning *pyr-Gal4*: GCCAAGCTTGCATGCCTCTAGA TGACATTCTGCAGATACGGGTAGTTC | This paper | N/A |
| Primer used for *pyr-Gal4* screen: GATCTCACGATCGGCCGTAAAATG | This paper | gRNAseqF2 |
| Primer used for *pyr-Gal4* screen: GATTTCGATTACACACACTCAATCTCTCG | This paper | gRNASeqR1 |
| Primer used for *pyr-Gal4* screen: GGATGCTATTAACCCTGAACTTTC | This paper | pJet seqR |
| Primer used for *pyr-Gal4* screen: CGCAGGGGATTTCTCC | This paper | CseqF |
| Primer used for *pyr-Gal4* screen: CCAGATTGAAATCGCG | This paper | Seq2F |
| Primer used for *pyr-Gal4* screen: CCAATGGCTAATATGCAG | This paper | Seq2R |
| gRNA for *pyr-Gal4*: ATAATATAAGTCCTGACATTGGG | This paper | N/A |
| Primer for cloning *htl-enh-FRT-stop-FRT3-FRT-FRT3-Gal4*: AATTCGAGCTCGGTACCGCTAGCGG CAAGGAGAAATTCCAACGCAGAGAC | This paper | N/A |
| Primer for cloning *htl-enh-FRT-stop-FRT3-FRT-FRT3-Gal4*: GATGAACGGGTGGGGATGG | This paper | N/A |
| Primer for cloning *htl-enh-FRT-stop-FRT3-FRT-FRT3-Gal4*: CGAGACCGGCACGAGTCTG | This paper | N/A |
| Primer for cloning *htl-enh-FRT-stop-FRT3-FRT-FRT3-Gal4*: GGGAGATTTAAGAGAGGTAGAGAATC | This paper | N/A |
| Primer for cloning *htl-enh-FRT-stop-FRT3-FRT-FRT3-Gal4*: TGTTCCAAATTGGTCCGCGTAGTCC | This paper | N/A |
| Primer for cloning *htl-enh-FRT-stop-FRT3-FRT-FRT3-Gal4*: CTACGCGGACCAATTTGGAACAACCAAACCG AAAGACTTAATTTATATTTATTTAATTAATTTTAATAA AAC | This paper | N/A |
| Primer for cloning *htl-enh-FRT-stop-FRT3-FRT-FRT3-Gal4*: GGCAAAAAAAAAACTGAGAATTTGC | This paper | N/A |
| Primer for cloning *htl-enh-FRT-stop-FRT3-FRT-FRT3-Gal4*: GCCAAGCTTGCATGCCCTCCTCTATGCCTGAACCC AGC | This paper | N/A |

| | | |
|---|---|---|
| Primer for cloning *htl-LexA*: CACCGCAAGGAGAAATTCCAACGCAGAGAC | This paper | N/A |
| Primer for cloning *htl-LexA*: GCCAAGCTTGGCGAATTCTGTTCCAAATTGGTCCG CGTAGTCC | This paper | N/A |
| Primer for cloning *UAS-Htl:mcherry:* AATTCGAGCTCGGTACCGAATTCATGGCTGCCGCC TGG | This paper | N/A |
| Primer for cloning *UAS-Htl:mcherry:* GGATCTGATCAAATTTGCCCACC | This paper | N/A |
| Primer for cloning *UAS-Htl:mcherry:* CTGACGAGCACATTCCTGGC | This paper | N/A |
| Primer for cloning *UAS-Htl:mcherry:* GCCAAGCTTGCATGCCCTCGAGAT AATTACACCACTTCTGCAGGTTGTCC | This paper | N/A |
| Primer for cloning *UAS-Pyr:GFP*: AATTCGAGCTCGGTACCGCGGCCG CATGTTCCACAAGTTCATGCCCAATG | This paper | N/A |
| Primer for cloning *UAS-Pyr:GFP*: AGCTCCTCGCCCTTGGACATG GTTGTTGTGGTTGTTGTTGTTGTG | This paper | N/A |
| Primer for cloning *UAS-Pyr:GFP*: CAACAACAACCACAACAACCATGTCCAAGGGCGA GGAGC | This paper | N/A |
| Primer for cloning *UAS-Pyr:GFP*: GCCAAGCTTGCATGCGCTAGC TGGTGTCTTGTACAGCTCATCCATGCCC | This paper | N/A |
| Primer for cloning *UAS-Pyr:GFP*: GGCATGGATGAGCTGTACAAGACACCAGCTAGCC CAGTGG | This paper | N/A |
| Primer for cloning *UAS-Pyr:GFP*: GCCAAGCTTGCATGCCGGATCCCTCGAGCTATAA ATCTATATAATACAAGCTAACAAAATACTTACCAC | This paper | N/A |
| Primer for cloning *UAS-Ths:GFP*: AATTCGAGCTCGGTACCGCGGCCGCAT GTCGAATCAGTTAGAGAGACTGCTG | This paper | N/A |
| Primer for cloning *UAS-Ths:GFP*: GCCGCCTTGCCCCTCGACGCTCTTCTTGGGCCCC ACAG | This paper | N/A |
| Primer for cloning *UAS-Ths:GFP*: GTCGAGGGGCAAGGCGGCATGTCCAAGGGCGAG GAGC | This paper | N/A |
| Primer for cloning *UAS-Ths:GFP*: ATGTCCAAGGGCGAGGAGC | This paper | N/A |
| Primer for cloning *UAS-Ths:GFP*: ACTGCCGCCGCCACTGCCCTTGTACAGCTCATCCA TGCCC | This paper | N/A |
| Primer for cloning *UAS-Ths:GFP*: GGCAGTGGCGGCGGCAGTGTCAACGATGCCTGCT ACATGTTC | This paper | N/A |
| Primer for cloning *UAS-Ths:GFP*: GCCAAGCTTGCATGCCTCTAGAC TACGCAAATCTCTGATGAGTGAACC | This paper | N/A |
| Recombinant DNA | | |
| pUC19 | Addgene | 50005 |

| | | |
|---|---|---|
| pUASt | DGRC | 1000 |
| pACT-FRT-stop-FRT3-FRT-FRT3-Gal4 | Addgene | 52889 |
| pBPnlsLexA::p65Uw | Addgene | 26230 |
| pCFD3 | [6] | N/A |
| pCFD3-pyr-Gal4-gRNA | This paper | N/A |
| pJet1.2-pyr-T2A-Gal4 | This paper | N/A |
| pLot-Htl:mCherry | This paper | N/A |
| pHtl-enh-FRT-stop-FRT3-FRT-FRT3-Gal4 | This paper | N/A |
| pBP-htl-enh-nlsLexA::p65Uw | This paper | N/A |
| p{GMR93H07-Gal4} | [7] | N/A |
| Software and Algorithms | | |
| Fiji- ImageJ 1.52p | ImageJ | https://fiji.sc |
| Prism 8.0 | GraphPad | https://www.graphpad.com/ |
| Adobe Photoshop 22.5.1 | Adobe | https://www.adobe.com |
| Adobe Illustrator 25.4.1 | Adobe | https://www.adobe.com |
| Microsoft Excel (Version 2111) | Microsoft | https://www.office.com |
| SnapGene 3.3.4 | SnapGene | https://www.snapgene.com |
| VassarStats | | vassarstats.net |
| Imaris 9.5.0 | Imaris | https://imaris.oxinst.com |
| Matlab 2019b | MathWorks | https://mathworks.com |
| Andor iQ3 | Oxford Instruments | https://andor.oxinst.com/products/iq-live-cell-imaging-software/ |
| Zen 3 | Carl Zeiss Microscopy GmbH | https://www.zeiss.com/microscopy/int/home.html?vaURL=www.zeiss.com/microscopy |
