## Supplementary material for "Cytonemes coordinate asymmetric signaling and organization in the *Drosophila* muscle progenitor niche": Movie Legends

### SUPPLEMENTARY MOVIES

**Supplementary Movie 1. 3D-rendered view of orthogonal AMP cytonemes visualized using live imaging.** The image was captured from the tip of the cytonemes to the ventral sides of the AMPs. Genotype: *htl-LexA, LexO-CD2:GFP*. Scalebar: 10μm (between two long ruler marks)

**Supplementary Movie 2. A time-lapse movie showing dynamics of AMP cytonemes in *ex vivo* cultured wing discs.** The time-lapse interval is 1 min. Genotype: *htl-LexA, LexO-CD2:GFP*. Scale bar: 5μm (between two long ruler marks).

**Supplementary Movie 3. Time-lapse movie showing growth of AMP cytonemes.** The time-lapse interval is 1 min. Genotype: *htl-LexA, LexO-CD2:GFP*. Scale bar: 5μm (between two long ruler marks).

**Supplementary Movie 4. Dynamic niche-sharing by multiple AMP cytonemes over time.** Time-lapse (1 min intervals) movie of a developing wing disc in ex vivo culture showing dynamic niche sharing by AMP cytonemes. Genotype: *htl-LexA, LexO-CD2:GFP*.

**Supplementary Movie 5. 3D-reconstruction of a wing disc showing the organization of orthogonal AMP cytonemes within the disc epithelium.** The 3D image was captured using a triple-view confocal microscope and processed as described in Methods. Genotype: *ths-Gal4/ UAS-nls:mCherry; htl-LexA/ LexO-CD2:GFP*. Color channels: AMP cytonemes (red) disc cell nuclei (blue) are pseudo-colored.

**Supplementary Movie 6. Serial YZ-stacks, showing AMP cytonemes (green) invading through the basolateral intercellular space between *ths*-expressing disc cells (red).** The nanoscopic resolution was achieved using Airyscan microscopy. Genotype: *ths-Gal4, UAS-mCherryCAAX/+ ; htl-LexA, LexO-CD2:GFP/+*. Scale bar: 2μm.

**Supplementary Movie 7. Synaptic contacts between AMP cytonemes and adherens junctions of the wing disc epithelium.** A 3D rendered view showing AMP cytonemes (green) grew between the intercellular space of wing disc cells (unmarked) and contacted Dlg-marked apical adherens junction of the wing disc epithelium (blue). The gap between the two Dlg-stained membranes represents the luminal space of the sac-like wing disc epithelium. Genotype: *ths-Gal4, UAS-mCherryCAAX/+; htl-LexA, LexO-CD2:GFP/+*. Scale bar: 1μm.

**Supplementary Movie 8. Endogenous Htl:GFP$^{fTRG}$ on niche-occupying AMP cytonemes.** 3D projection of image volume showing disc-invading AMP cytonemes localizing endogenous Htl:GFP$^{fTRG}$ puncta. Wing disc unmarked and on the left of marked AMPs. Both wing disc hinge and notum areas are shown. Genotype: *Htl:GFP$^{fTRG}$*. Scale bar: 20μm.
